# Transcriptomic analysis of aging mouse sciatic nerve reveals early pathways leading to sarcopenia

**DOI:** 10.1101/2022.02.01.478571

**Authors:** Nicole Comfort, Meethila Gade, Madeleine Strait, Samantha J. Merwin, Daphne Antoniou, Anna Memou, Hardy J. Rideout, Stefania Corti, Shingo Kariya, Diane B. Re

## Abstract

**Background:** Sarcopenia, the age-associated decline in skeletal muscle mass and strength, has long been considered a disease of muscle only, but accumulating evidence suggests that sarcopenia could originate from the neural components controlling muscles. To identify early molecular changes in the efferent nerves that may drive sarcopenia initiation, we performed a longitudinal transcriptomic analysis of the sciatic nerve in aging mice.

**Methods:** Sciatic nerve and gastrocnemius muscle were obtained from young adult, middleaged, old, and sarcopenic (5,18, 21 and 24 months old, respectively) C57BL/6J female mice (n=6 per age group). Sciatic nerve RNA was extracted and subjected to RNA sequencing (RNA-seq), with real-time quantitative reverse transcription PCR (qRT-PCR) validation of differentially expressed genes (DEGs). Functional enrichment analysis of clusters of genes associated with patterns of gene expression across age groups was performed. Sarcopenia was confirmed with qRT-PCR of previously established markers of sarcopenia onset in gastrocnemius muscle.

**Results:** We detected 33 significant DEGs in sciatic nerve of 18-month-old mice compared to 5-month-old mice (absolute value of fold change > 2; false discovery rate [FDR] < 0.05) which we validated with qRT-PCR of the three top up- and down-regulated genes. Up-regulated genes were associated with circadian rhythm and the AMPK signaling pathway, while down-regulated genes were associated with biosynthesis and metabolic pathways and circadian rhythm. Strikingly, we detected a significant increase in *Myog* expression (log_2_ fold change = 18.93, FDR *q*-value = 1.54×10^−12^) in sciatic nerve of 18-month-old mice, before up-regulation in muscle was observed. We identified seven clusters of genes with similar expression patterns across groups. Functional enrichment analysis of these clusters revealed biological processes that may be implicated in sarcopenia initiation including extracellular matrix organization and circadian regulation of gene expression.

**Conclusions:** Gene expression changes in mouse peripheral nerve can be detected prior to overt clinical onset of sarcopenia. These early molecular changes we report shed a new light on biological processes that may be implicated in sarcopenia initiation and pathogenesis. Future studies will validate which of the key changes we reported have disease modifying and/or biomarker potential.

## 1. Introduction

Based on its symptomatic and histological features, sarcopenia, the age-associated loss in skeletal muscle mass and strength, has long been considered a disease of skeletal muscle fibers only. However, accumulating evidence suggests that sarcopenia could originate from the neural components controlling muscle strength and hence be defined as a neuromuscular disease (Krishnan et al. 2016; Pannérec et al. 2016; Rolland et al. 2008; Ryall, Schertzer, and Lynch 2008). Although current evidence suggests that motor nerves and presynapses of neuromuscular junctions (NMJs) could be implicated in sarcopenia pathophysiology, their role in initiation of sarcopenia has not been fully elucidated.

We hypothesized that sarcopenia onset and its early molecular programming could be captured in the major nerve of the lower limb, the sciatic nerve, reflected by changes in gene expression in nerves preceding sarcopenia development in muscle. Most previous gene expression profiling studies investigating the mechanisms of sarcopenia pathophysiology have been limited by their use of microarrays for peripheral nerve molecular profiling. Microarrays cover only a defined set of transcripts, have high background levels due to cross-hybridization, and have a limited dynamic range due to both background as well as saturation of signals. However, studies that utilized next-generation sequencing technologies have so far focused on age-related changes in muscles rather than nerves (Zhou et al. 2018).

To circumvent these issues and capture early molecular events responsible for the initiation of sarcopenia, we performed deep RNA sequencing (RNA-seq) (Wang, Gerstein, and Snyder 2009) of sciatic nerves, isolated at different time points from healthy young (5 months), middle-aged (18 and 21 months), and sarcopenic (24 months) old mice. Our objective was to identify early changes in gene expression related to the development of sarcopenia via transcriptomic profiling of aging sciatic nerves *in vivo*. In agreement with a potential neurogenic origin of sarcopenia, we identified a series of relevant gene expression changes in mouse peripheral nerve prior to overt clinical onset of sarcopenia in muscles. Among the early molecular events that we report at 18 months, we confirm the importance of biological processes previously linked to sarcopenia while also reporting new findings that shed light on other pathophysiological mechanisms that may be implicated in sarcopenia initiation and pathogenesis. To our knowledge, this study is the first to utilize untargeted RNA-seq to investigate the transcriptome of sciatic nerves in the context of sarcopenia. Future studies are warranted to confirm whether neuromuscular biomarkers of sarcopenia could be identified in efferent nerves at even earlier timepoints and to validate which among them are phenotypic drivers with potential implications for sarcopenia prevention and therapy.

## 2. Methods

### 2.1. Mice and sample collection

Female C57BL/6J mice were obtained from the National Institute on Aging mouse colony, two weeks before turning 5, 18, 21, and 24 months old. The mice were kept in the Columbia University Medical Center animal facility for two weeks and then were used immediately. The mice had free access to the standard pellet and water diet, and were maintained under a constant 12-hour light/dark cycle at an environmental temperature of 21–23 °C. All animal procedures complied with the Guide for the Care and Use of Laboratory Animals and were approved by the Columbia University Institutional Animal Care and Use Committee (Approval ID: AC-AAAN8900).

At 5 (young adult negative control), 18 (middle-aged), 21 (old), and 24 (sarcopenic positive control) months, mice (n=6 per group) were euthanized via deep anesthesia followed by decapitation. Sciatic nerves were quickly dissected before undergoing RNA extraction. The remaining leg was snap-frozen in liquid nitrogen and stored at −80°C.

### 2.2. RNA extraction

#### 2.2.1. Sciatic nerve

Total RNA was isolated from freshly dissected sciatic nerves using the Invitrogen PureLink™ RNA Mini Kit (Cat # 12183018A) following the manufacturer’s protocol for ≤ 10 mg fresh soft animal tissue. All samples had total RNA concentration and RNA integrity numbers (RINs) assessed prior to sequencing or real-time quantitative reverse transcription PCR (qRT-PCR). RINs were evaluated using an Agilent 2100 Bioanalyzer and ranged from 6.2 to 9.6 (24 samples; average: 8.7, **Supplemental Table 1**).

#### 2.2.2. Gastrocnemius muscle

The gastrocnemius muscles were dissected from the leg on dry ice and 50 mg of tissue was flash frozen in duplicate. The tissue samples were then placed in a pre-chilled Eppendorf tube and homogenized with a Qiagen TissueRuptor II (Cat # 9002755) with disposable probes (Qiagen, Cat # 990890). Total RNA extraction was performed using trizol-chloroform extraction followed by column centrifugation using the Invitrogen PureLink™ RNA Mini Kit following the manufacturer’s instructions. Then, samples were treated with DNAse followed by wash buffer and eluted in water. Total RNA was quantified using the RiboGreen™ RNA Quantitation Kit (Thermo Fisher, Cat # R11490).

### 2.3. Library preparation and RNA sequencing

Libraries were constructed at the Columbia Biomedical Core Facility. Briefly, mRNA was enriched from total RNA via poly-A pull-down using the Illumina TruSeq Stranded mRNA kit according to the manufacturer’s instructions. Libraries were 2×100 base-pair (bp) paired-end sequenced for each sample on an Illumina HiSeq4000 using 75-cycle High Output Kits (target 60 million reads per sample) at the Columbia University Genome Center. Raw sequencing data and processed data will be deposited at the NIH Gene Expression Omnibus (GEO).

### 2.4. RNA sequencing analysis, quality control, and identification of differentially expressed genes

We used RTA (Illumina) for base calling and bcl2fastq2 (version 2.17) for converting BCL to fastq format, coupled with adaptor trimming. Quality control of the raw and trimmed reads was performed using FastQC and MultiQC (Andrews 2010; Ewels et al. 2016). We aligned trimmed reads that passed quality control (Phred score > 30) to the mouse reference genome and transcriptome annotation (mm10, UCSC) using STAR aligner (version 2.5.2b) (Dobin and Gingeras 2015) and quantified the reads that aligned uniquely to the transcriptome using featureCounts (v1.5.0-p3) (Liao, Smyth, and Shi 2014). Sequencing yielded libraries with an average size of 39 million reads after alignment.

We used the DESeq2 (version 1.34.0) package for R (version 4.1.1; R-project.org/) (Love, Huber, and Anders 2014) for further quality control, exploratory data anslysis, and differential expression analysis. For sample-level quality control, we conducted principal component analysis (PCA) of the RNA-seq data by first normalizing the counts for each sample using size factors obtained from DESeq2, log-transforming the normalized counts (using the regularized-logarithm [rlog] transformation), and computing the principal components of the resulting matrix using the plotPCA function of DESeq2 (**Supplemental Figure 1A**). We also created heatmaps of the correlation of rlog-transformed gene expression for all pairwise combinations of samples (**Supplemental Figure 1B**). Based on this quality control assessment, we removed two samples that were outliers from downstream analyses (resulting PCA plot and correlation heatmap of rlog-transformed gene expression shown for *N*=22 shown in **Supplemental Figure 2**).

We removed 4,749 (20.3%) rows of the DESeqDataSet that had no counts or only a single count across all included samples (*N*=22). We additionally specified that at least 20% of samples (i.e., at least 5 samples) have a count of 10 or higher, resulting in removal of an additional 3,079 genes (13.1%). In total, 7,828 (33.4%) of 23,416 unique genes were filtered out, resulting in 15,588 genes analyzed in the DESeqDataSet. DESeq2 fits negative binomial generalized linear models for each gene and determines differential expression for pairwise comparisons by applying the Wald test for significance testing. Adjusted *p*-values were calculated by Benjamini-Hochberg false discovery rate (FDR). We considered genes that had an absolute value of a log_2_FoldChange (LFC) > 1.0 with adjusted p-value < 0.05 to be differentially expressed genes (DEGs).

### 2.5. Validation of differentially expressed genes with qRT-PCR

#### 2.5.1. SYBR® Green RT-PCR

For select genes in sciatic nerve identified as differentially expressed by DESeq2, we validated the RNA-seq results using qRT-PCR. We also performed qRT-PCR of the genes *Chrnd, Gadd45a, Myog, Runx1, Chrng*, and *Ncam* in gastrocnemius muscle to confirm onset of sarcopenia. For qRT-PCR, Qiagen QuantiTect Primers were reconstituted in TE buffer (pH 8.0). Primer details are listed in **Supplemental Table 2**. The reverse transcription reaction was performed using the Promega GoScript™ Reverse Transcriptase kit (Cat # PRA5000) for a reaction volume of 20 μL. This was followed by SYBR^®^ Green RT-qPCR reaction using KiCqStart™ SYBR^®^ Green qPCR ReadyMix™ (Millipore Sigma, Cat # KCQS00). Briefly, 5 μL reaction mix was prepared using 2.5 μL qPCR ready mix, 1.25 μL sample cDNA, and 1.25 μL primers. The RT-PCR reaction was run on Bio-Rad CFX384 Touch Real-Time PCR System as per kit instructions for three technical replicates.

#### 2.5.2. qRT-PCR data analysis

RT-PCR results were analyzed in RStudio (version 3.6.0). Data was cleaned by excluding samples with cycle threshold (Ct) values > 35 and only 1 read/technical replicate. Raw Ct values were then normalized to the geometric mean of housekeeping genes (*Ppia* and *Hprt*, **Supplemental Figure 3**). The resulting delta Ct (dCt) values were further normalized to the 5-month age group, which served as the reference, to create delta-delta Ct values (ddCT). Plots were generated using GraphPad Prism (version 8.4.0). Only the raw dCT values were plotted for assays (*Chrng, Myog, Mstn*) with many non-detects in the reference group. A one-way ANOVA was used to test for any statistically significant differences between the groups followed by post hoc analysis using Fisher’s Least Significant Difference (LSD) test. We report significant differences in ddCT comparing each group (18, 21, 24 months) to the reference group (5 months).

### 2.6. Pattern identification

We analyzed all pair-wise comparisons simultaneously using the Likelihood Ratio Test (LRT) in DESeq2 to identify any genes that showed an interesting change in expression across the different groups (5, 18, 21, 24 months). We subset the rlog-transformed normalized counts to contain only genes significant with adjusted *p*-value < 0.05 and explored clusters of genes with similar expression patterns across sample groups using the ‘degPatterns’ function from the DEGreport R package. Gene lists associated with each identified cluster were used as input for functional enrichment analysis.

### 2.7. Functional and pathway enrichment analysis

DEGs identified by the Wald test were explored using gene ontology (GO) functional enrichment analysis and KEGG (Kyoto Encyclopedia of Genes and Genomes) pathway enrichment analysis performed using the database for annotation, visualization and integrated discovery (DAVID) (https://david.ncifcrf.gov/) (Sherman et al. 2007; Dennis et al. 2003; Huang et al. 2007; Huang, Sherman, and Lempicki 2009). Pathways or GO terms were considered significant at *p*-values < 0.05.

To investigate the biological relevance of clusters of DEGs associated with main expression patterns, we performed GO functional enrichment analysis using GOrilla, a publicly available GO enrichment tool (http://cbl-gorilla.cs.technion.ac.il) (Eden et al. 2009). GOrilla compares a target gene list (in this case, genes assigned to a given cluster) to a background set of genes (*N*=2,925 genes significant in the LRT with FDR < 0.05) and assesses the significance of enrichment for GO terms using the hypergeometric test.

## 3. Results

### 3.1. Confirmation of sarcopenia onset in C57BL/6J mice via qRT-PCR of genes in gastrocnemius muscle

To validate the onset of overt sarcopenia in our different aged mouse groups in an unbiased manner (independent of mouse size/weight/basal activity/hydration, etc.), we performed qRT-PCR quantification of the following genes, associated with NMJ denervation, in gastrocnemius muscle: nicotinic acetylcholine receptor (nAChR) gamma and delta subunits (*Chrng* and *Chrnd*), runt-related transcription factor-1 (*Runx1*), growth arrest and DNA damageinducible 45α (*Gadd45a*), and Myogenin (*Myog*) It has been previously established that increased expression of these genes in aging muscle is concomitant with the transition to sarcopenia in 24-month-old female C57BL/6J mice. As reported previously (Barns et al. 2014), we observed significantly increased expression in *Chrnd, Chrng, Runx1*, and *Gadd45a* at age 24 months (**Supplemental Figure 4**). Note that for *Chrng*, the substantial increase in expression measured at 24 months was not tested for statistical significance because of the very low expression (mostly non-detectable) in the reference group (5 m old). There was an increase in *Myog* expression at 24 months that was not statistically significant.

### 3.2. Identification of differentially expressed genes (DEGs) in sciatic nerve by RNA-seq

The number of DEGs (|LFC| > 1.0, FDR < 0.05) identified using DESeq2 is shown in **Table 1.** The full DESeq2 results, including the list of these DEGs complete with their full gene name, associated LFC and *p*-values, are shown in **Supplemental Spreadsheet 1.** Because we were interested in the earliest transcriptional changes that may precede initiation of sarcopenia pathology, we assessed the contrast between 18-month-old and 5-month-old mice in more detail. Comparing these two groups, a total of 33 DEGs were identified, including 16 up-regulated and 17 down-regulated DEGs (**Figure 1**, **Table 2**). Interestingly, the most up-regulated gene (according to LFC) was *Myog* with a LFC = 18.93 (FDR *q*-value = 1.54×10^−12^). Volcano plots of the other pairwise comparisons are shown in **Supplemental Figure 5**.

**Figure 1.**
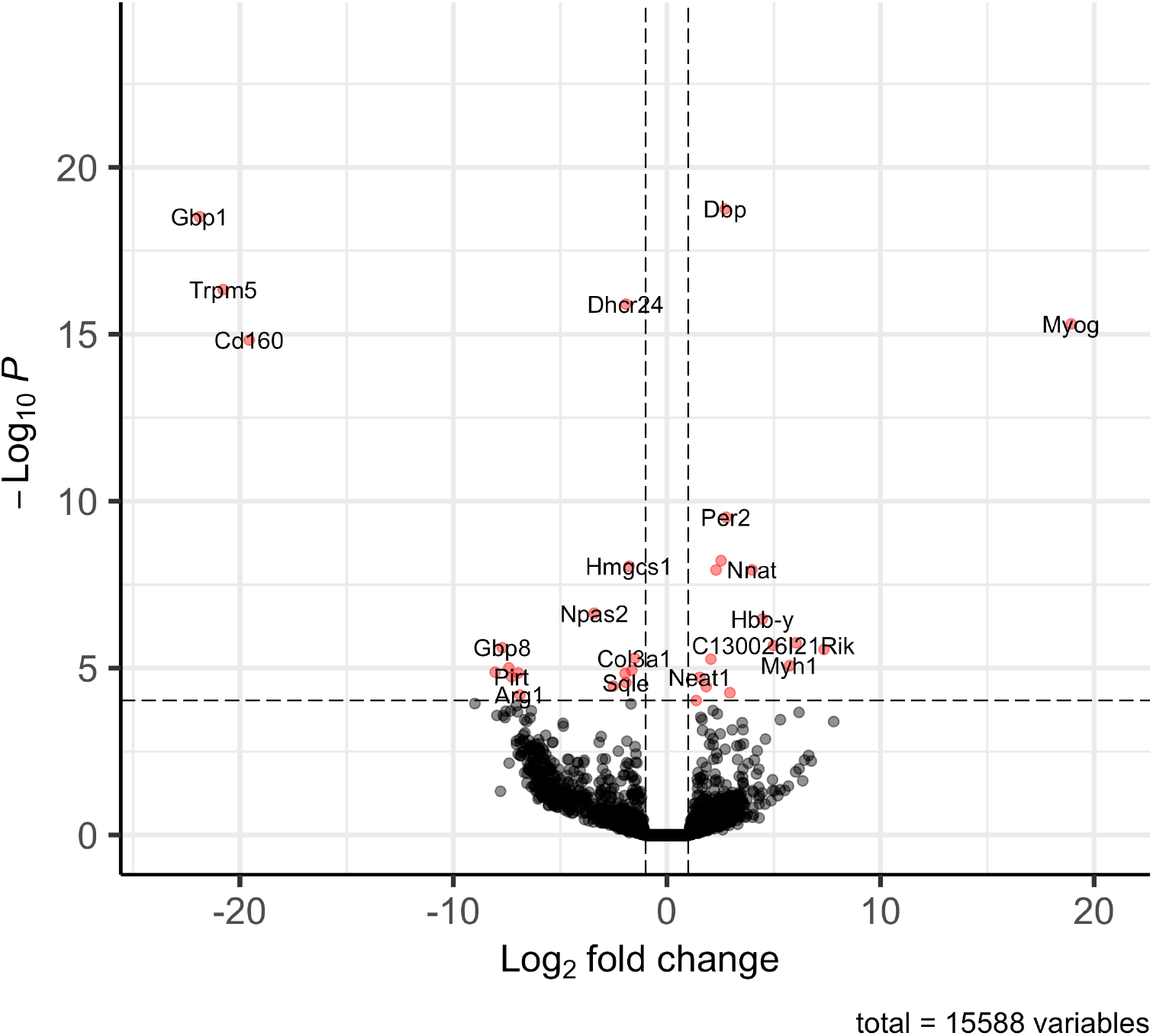
Volcano plot of differentially expressed genes (of 15,588 genes analyzed) in sciatic nerve comparing 18-month-old mice (n=5) to 5-month-old mice (n=6). The dashed line is the FDR q-value = 0.05. The 33 genes significant at |LFC| > 1.0 and FDR q-value < 0.05 are plotted in red.

**Table 1.** Number (%) of differentially expressed genes (DEGs) out of 15,588 genes.

| # DEGs FDR < 0.05 |  |  |
| --- | --- | --- |
| Contrast | LFC > 1.0 | LFC < -1.0 |
| 18 vs 5 months | 16 (0.1%) | 17 (0.11%) |
| 21 vs 5 months | 18 (0.12%) | 10 (0.064%) |
| 24 vs 5 months | 50 (0.32%) | 18 (0.12%) |
| 21 vs 18 months | 13 (0.083%) | 5 (0.032%) |
| 24 vs 18 months | 30 (0.19%) | 1 (0.0064%) |
| 24 vs 21 months | 3 (0.019%) | 1 (0.0064%) |

**Table 2.** The 33 DEGs identified in sciatic nerve comparing 18-month-old mice (n=5) to healthy, young control mice 5 months old (n=6), including 16 up-regulated genes and 17 down-regulated genes.

|  | Gene | Log<sub>2</sub> Fold Change | FDR *q*-value |
| --- | --- | --- | --- |
| Upregulated | *Myog* | 18.93 | 1.54E-12 |
|  | *Lmod2* | 7.35 | 2.51E-03 |
|  | *Hba-x* | 6.04 | 1.96E-03 |
|  | *Myh1* | 5.74 | 6.56E-03 |
|  | *C130026I21Rik* | 4.98 | 2.19E-03 |
|  | *Hbb-y* | 4.47 | 4.05E-04 |
|  | *Nnat* | 3.98 | 1.62E-05 |
|  | *Lep* | 2.95 | 2.76E-02 |
|  | *Per2* | 2.77 | 6.72E-07 |
|  | *Dbp* | 2.72 | 2.37E-15 |
|  | *Ppp2r2c* | 2.54 | 1.18E-05 |
|  | *Col24a1* | 2.30 | 1.62E-05 |
|  | *Tmem45b* | 2.06 | 4.42E-03 |
|  | *Per3* | 1.84 | 1.86E-02 |
|  | *Neat1* | 1.54 | 1.09E-02 |
|  | *Tef* | 1.37 | 4.40E-02 |
| Downregulated | *Col3a1* | -1.52 | 4.42E-03 |
|  | *Hmgcr* | -1.65 | 8.32E-03 |
|  | *Hmgcs1* | -1.78 | 1.54E-05 |
|  | *Sqle* | -1.92 | 1.51E-02 |
|  | *Dhcr24* | -1.93 | 4.87E-13 |
|  | *Arntl* | -1.96 | 8.97E-03 |
|  | *Mvd* | -2.52 | 1.84E-02 |
|  | *Npas2* | -3.40 | 2.98E-04 |
|  | *Arg1* | -6.88 | 3.12E-02 |
|  | *P2rx1* | -6.96 | 8.97E-03 |
|  | *Pirt* | -7.28 | 1.08E-02 |
|  | *Tnip3* | -7.40 | 7.32E-03 |
|  | *Gbp8* | -7.71 | 2.38E-03 |
|  | *Adamts16* | -8.05 | 8.96E-03 |
|  | *Cd160* | -19.56 | 3.90E-12 |
|  | *Trpm5* | -20.78 | 2.39E-13 |
|  | *Gbp1* | -21.89 | 2.37E-15 |

To explore the biological roles of up- and down-regulated DEGs, we performed functional and pathway enrichment analyses on the DEGs comparing 18-month-old and 5-month-old mice (listed in Table 2) using DAVID. Up-regulated DEGs were associated with biological processes and KEGG pathways related to circadian rhythm (*p* = 0.02), circadian entrainment (*p*=0.06), and the AMP-activated protein kinase (AMPK) signaling pathway (*p*=0.08), while down-regulated DEGs were associated with various pathways related to biosynthetic processes, including sterol, cholesterol, steroid, and isoprenoid biosynthetic processes (**Table 3**, **Supplemental Table 3**). Circadian rhythm was also enriched (*p*=0.04) among down-regulated genes including neuronal PAS domain protein 2 (*Npas2*) and the gene encoding Aryl hydrocarbon receptor nuclear translocator-like protein 1 (*Arntl*), which both displayed similar gene expression patterns across age groups (**Supplemental Figure 6**).

**Table 3.** KEGG pathway analysis of up- and down-regulated DEGs comparing 18-month-old mice (n=5) to 5-month-old mice (n=6).

|  | KEGG Pathway | Gene count | P-value | FDR q-value | Genes |
| --- | --- | --- | --- | --- | --- |
| Upregulated | Circadian rhythm | 2 | 2.00E-02 | 4.60E-01 | *Per2, Per3* |
|  | Tight junction | 2 | 5.70E-02 | 4.60E-01 | *Myh1, Ppp2r2c* |
|  | Circadian entrainment | 2 | 6.20E-02 | 4.60E-01 | *Per2, Per3* |
|  | AMPK signaling pathway | 2 | 8.10E-02 | 4.60E-01 | *Lep, Ppp2r2c* |
| Downregulated | Biosynthesis of antibiotics | 5 | 1.10E-04 | 2.50E-03 | *Hmgcr, Hmgcs1, Arg1, Mvd, Sqle* |
|  | Terpenoid backbone biosynthesis | 3 | 3.80E-04 | 4.40E-03 | *Hmgcr, Hmgcs1, Mvd* |
|  | Metabolic pathways | 6 | 1.50E-02 | 1.10E-01 | *Dhcr24, Hmgcr, Hmgcs1, Arg1, Mvd, Sqle* |
|  | Steroid biosynthesis | 2 | 2.40E-02 | 1.40E-01 | *Dhcr24, Sqle* |
|  | Circadian rhythm | 2 | 4.00E-02 | 1.80E-01 | *Arntl, Npas2* |

### 3.3. Validation of DEGs in sciatic nerve via qRT-PCR

We performed qRT-PCR on a set of DEGs to validate the RNA-seq results. As shown in **Figure 2**, the gene expression in the sciatic nerve measured by qRT-PCR agreed with our RNA-seq findings. Compared to 5-month-old mice, there was a significant increase in expression of period circadian regulator 2 (*Per2*, *p*=0.0072), D site albumin promoter binding protein (*Dbp*, *p*=0.0039), and protein phosphatase 2, regulatory subunit B, gamma (*Ppp2r2c*, *p*=0.0004) (Figure 2A-C) in 18-month-old mice. We additionally confirmed that the genes guanylate binding protein 11 (*Gbp1*), 24-dehydrocholesterol reductase (*Dhcr24*), and 3-hydroxy-3-methylglutaryl-Coenzyme A synthase 1 (*Hmgcs1*) were significantly down-regulated in 18-month-old mice compared to 5-month-old mice (*p*=0.0009 for *Gbp1* and *p*<0.0001 for *Dhcr24* and *Hmgcs1*) (Figure 2D-F). We also confirmed the upregulation of *Myog* in 18-month-old mouse sciatic nerve compared to 5-month-old mice (**Figure 3**). *Myog* was not detected in sciatic nerve of 5-month-old mice.

**Figure 2.**
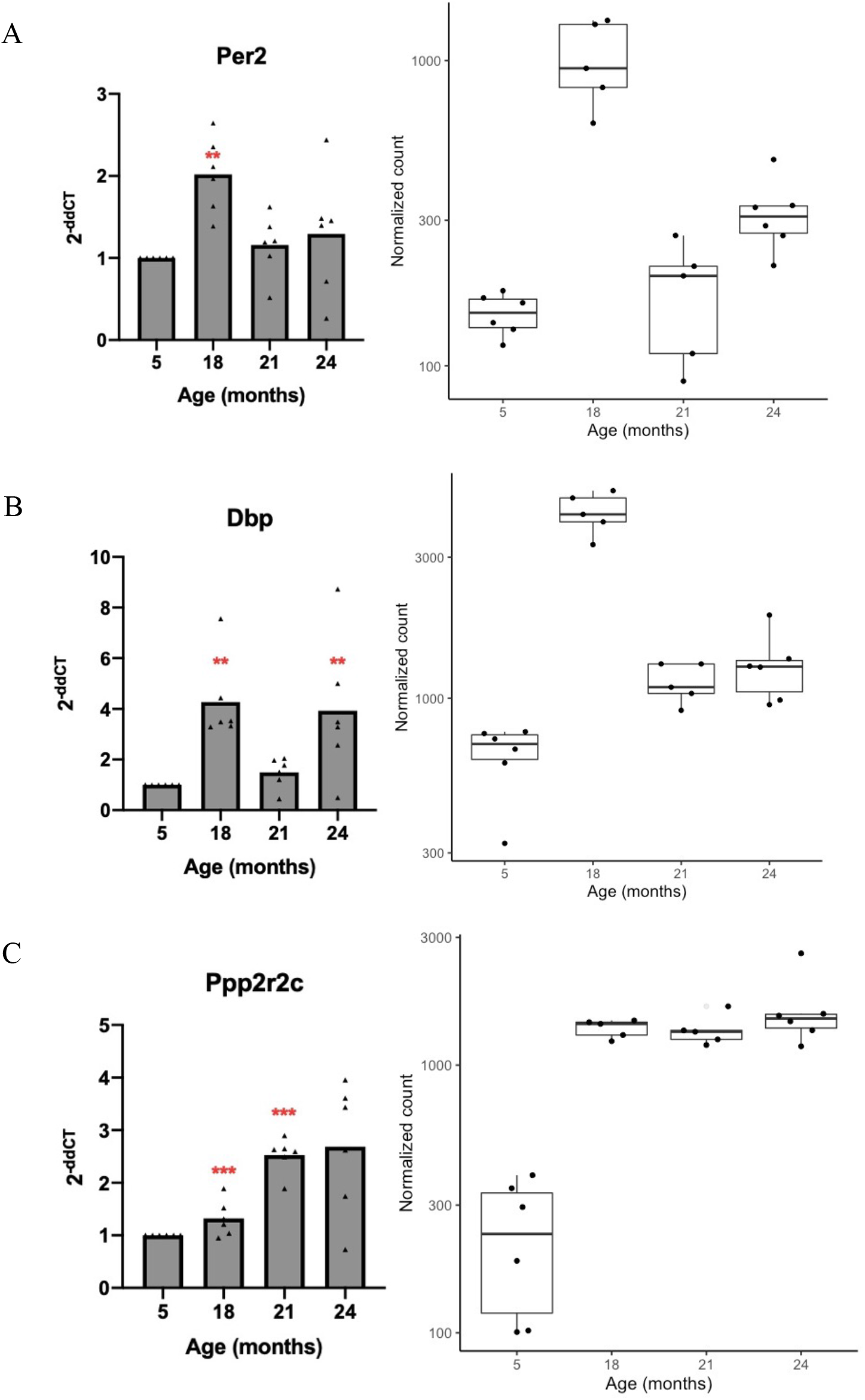
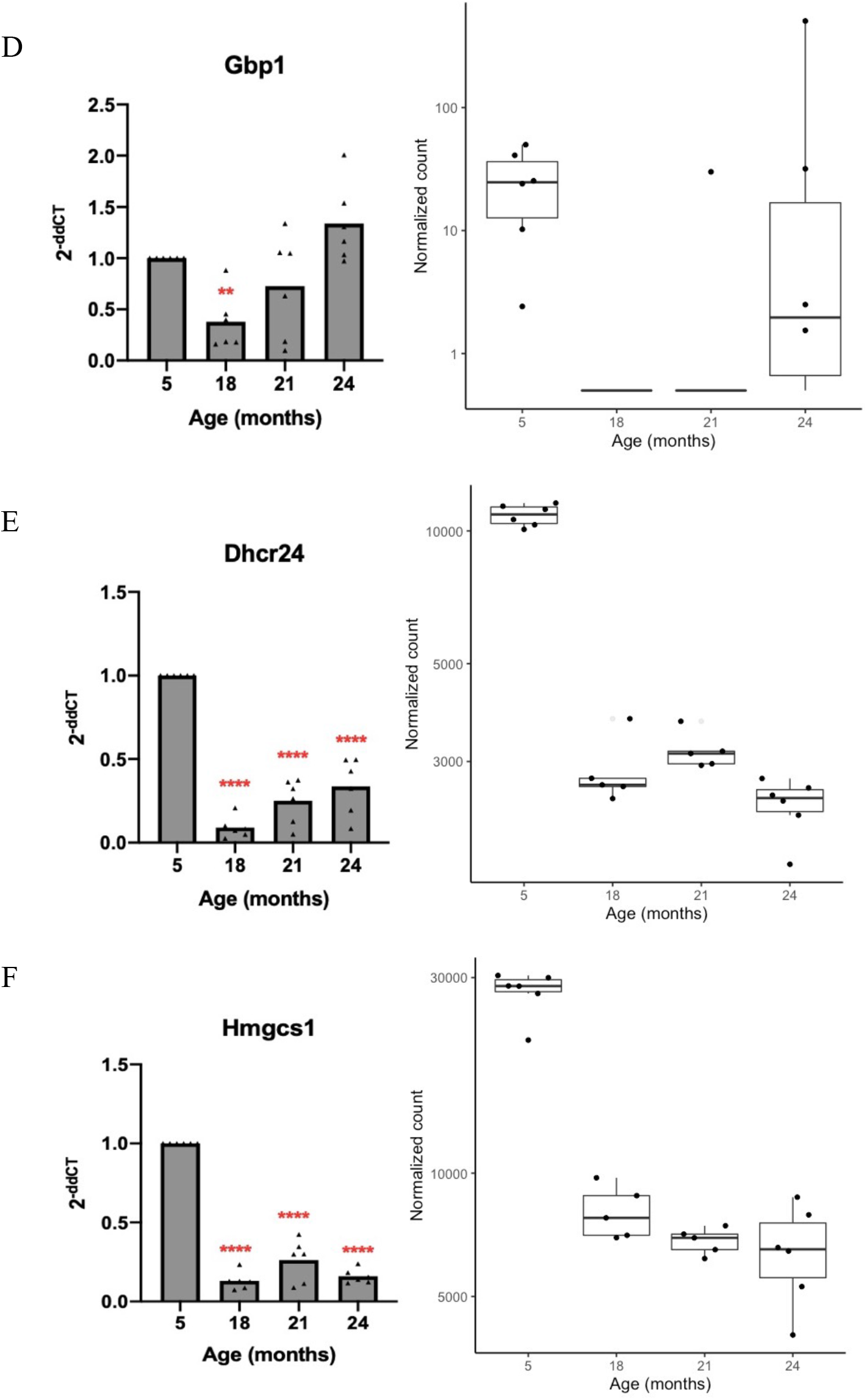
Gene expression of up-regulated (A-C) and down-regulated (D-F) genes in sciatic nerve of mice aged 5, 18, 21, and 24 months assessed by qRT-PCR and RNA-seq. Left: Relative gene expression assessed by qRT-PCR; Right: Corresponding plot of normalized counts from RNA-seq analysis (n=5-6 per age group). For RT-PCR, the delta-delta Ct was calculated using *Ppia* and *Hprt* housekeeping genes with 5-month-old mice used as the reference; n=5-6 mice per age group, except for the 5-month-old mice in (A) and (C) which had n=3 and 2, respectively, due to many samples being below the detection threshold. \*\**p* < 0.008, \*\*\**p* < 0.0005, \*\*\*\**p* < 0.0001, significant difference between group with reference group, ANOVA with Fisher’s LSD.

**Figure 3.**
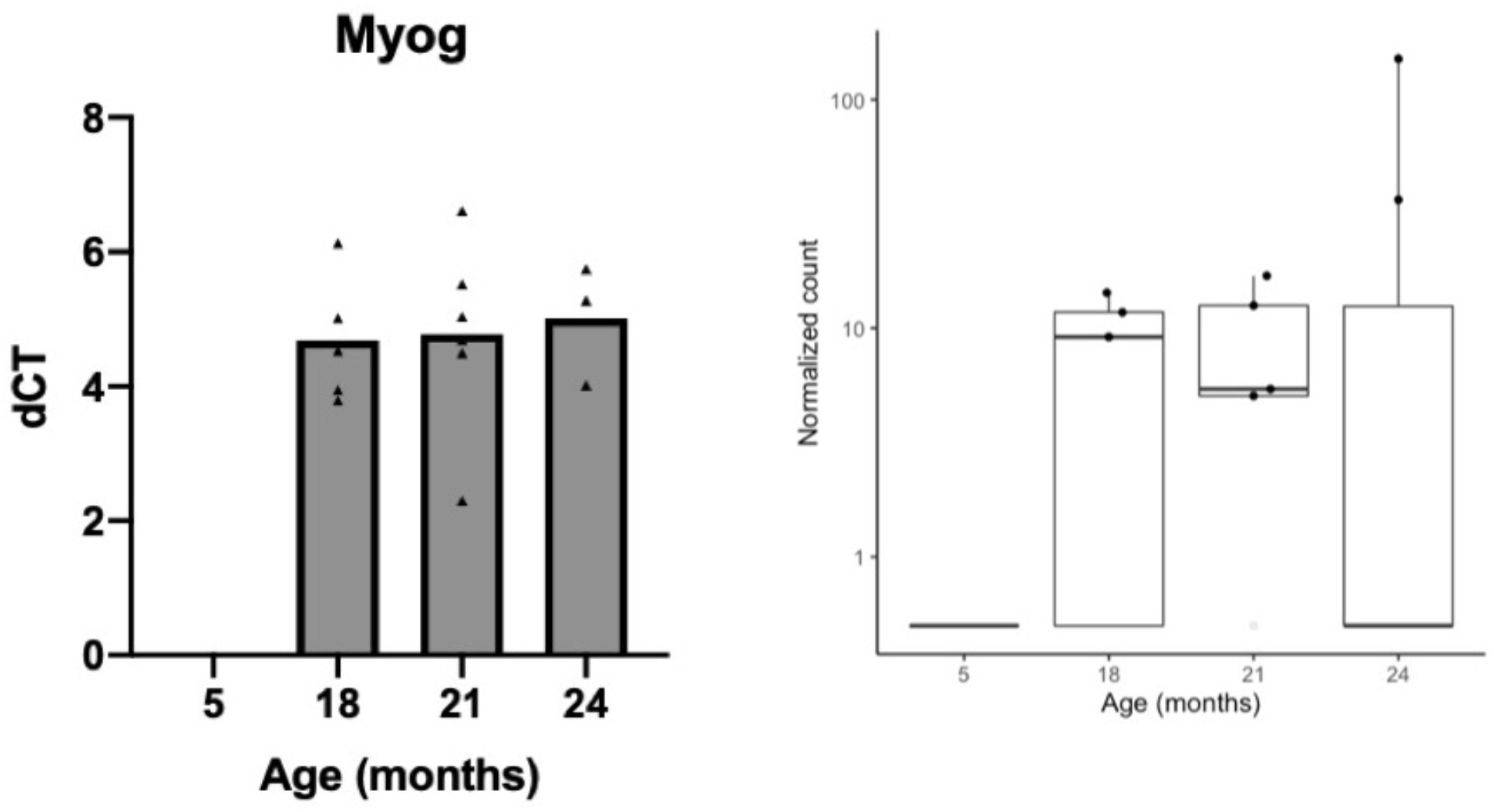
*Myogenin* (*Myog*) expression in sciatic nerve of mice aged 5, 18, 21, and 24 months. For the RT-PCR analysis, the number of samples for mice aged 5, 18, 21, and 24 months are n=0 (not detected), 5, 6, and 3, respectively.

### 3.4. Pattern identification and functional enrichment analyses of DEGs

To investigate biologically relevant changes in gene expression across age groups, we analyzed all pair-wise comparisons of the normalized RNA-seq counts simultaneously using the Likelihood Ratio Test. This test identified 2,925 genes significant at FDR < 0.05. We then identified clusters of genes within this list with similar expression patterns across age groups using the ‘degPatterns’ function in R. We identified seven clusters of genes containing 95-715 genes per cluster (**Figure 4**). Using GOrilla, we explored the biological relevance of these clusters using functional enrichment analysis assessing enrichment of clusters for GO Biological Process terms. The results of the functional enrichment analysis are shown in **Figure 5**.

**Figure 4.**
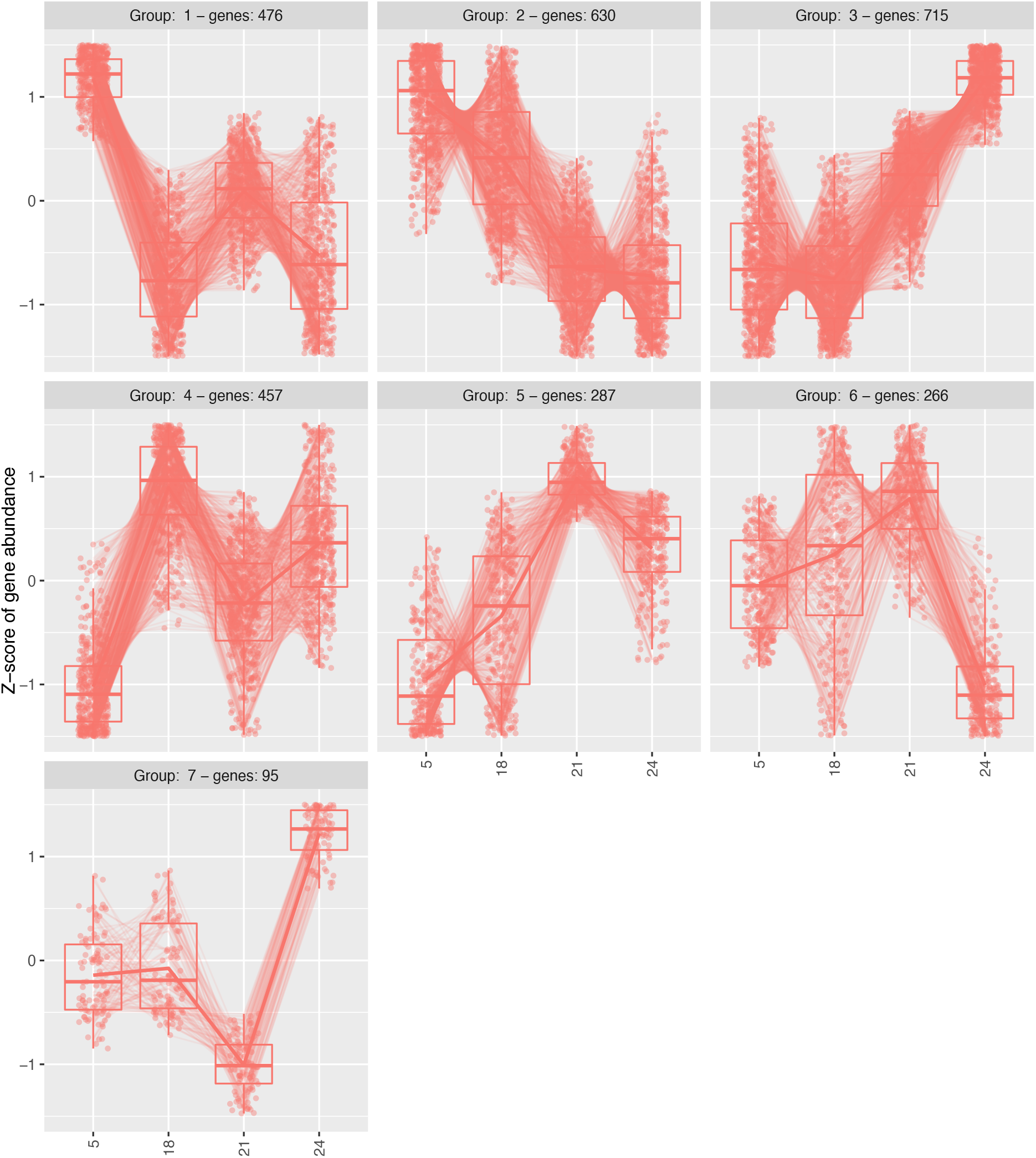
Clusters of genes with similar pattern of gene expression across age groups (labeled on the x axis). Identified from 2,925 genes significant in the Likelihood Ratio Test at FDR < 0.05.

**Figure 5.**
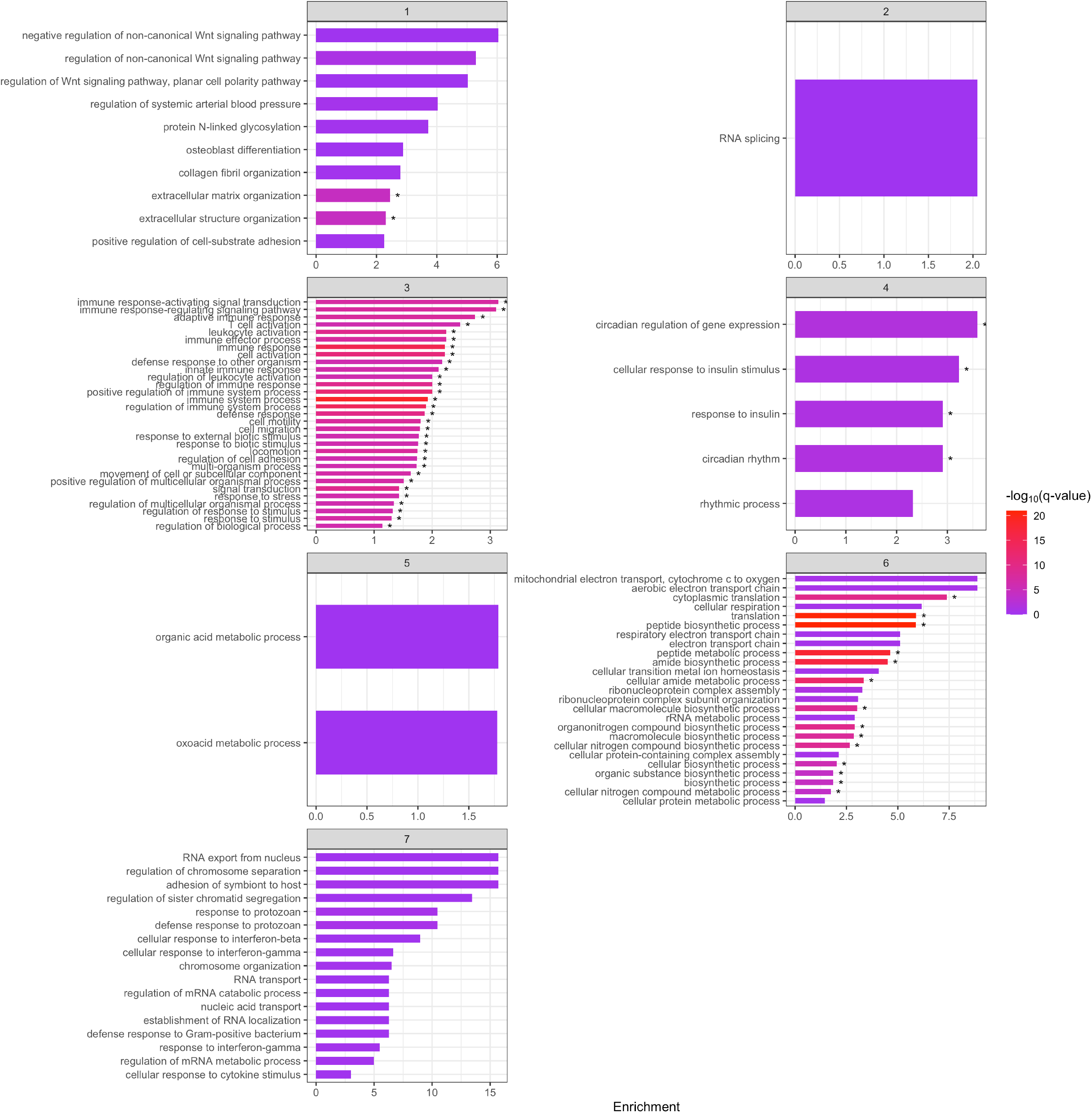
GO Biological Process functional enrichment analysis of the 7 clusters of gene expression, denoted in gray, identified by degPatterns. The Enrichment score is shown on the y axis, with bars colored according to −log_10_(*q*-value). *GO biological process terms significantly enriched with target genes, FDR *q*-value < 0.05.

Cluster 1 contained 476 genes and the gene list was significantly enriched with genes associated with extracellular matrix organization (FDR *q*-value = 4.07×10^−5^) and extracellular structure organization (FDR *q*-value = 4.77×10^−5^). Cluster 3 contained 715 genes which were significantly associated with 266 GO Biological Process terms at FDR < 0.05. The top 31 terms are shown in Figure 5 including immune system process (FDR *q*-value = 2.33×10^−19^), immune response (FDR *q*-value = 8.04×10^−16^), and regulation of immune system process (FDR *q*-value = 5.74×10^−15^). Circadian regulation of gene expression (FDR *q*-value = 0.0164), the cellular response to insulin stimulus (FDR *q*-value = 0.039), response to insulin (FDR *q*-value = 0.026), and circadian rhythm (FDR *q*-value = 0.047) were significantly enriched GO terms of the 457 genes in Cluster 4. The 266 genes in Cluster 6 were significantly associated with 14 Biological Process GO terms including peptide biosynthetic process (FDR *q*-value = 1.07×10^−21^), translation (FDR *q*-value = 1.89×10^−21^), and peptide metabolic process (FDR *q*-value = 1.24×10^−18^). The full results are available in **Supplemental Spreadsheet 2**.

## 4. Discussion

Sarcopenia is the age-associated progressive decline in skeletal muscle mass and strength (Rolland et al. 2008; Landi et al. 1999; Rosenberg 1989). Such compromised muscle function restricts mobility and daily activity, greatly reducing the quality of life and causing a loss of independence among affected individuals. Low skeletal muscle mass or strength is also a frequent cause of disability in the elderly and is associated with many adverse outcomes including increased risk of falls and related fractures, morbidity, frailty, and mortality, making it a significant public health issue (Janssen et al. 2004; Metter et al. 2002; Abellan van Kan 2009).

Sarcopenia has long been considered a disease of skeletal muscle fibers only but accumulating evidence suggests that sarcopenia may originate in the nervous system. For example, electrophysiological examinations of patients have observed a reduction in motor unit numbers (Vandervoot and Symons 2001; Doherty et al. 1993) as well as an increase in the motor unit size (Tieland, Trouwborst, and Clark 2018; Lexell 1995), an indication of motor neuron death and distal motor neuropathy, respectively, resulting in decreased muscle strength. Thus, the neurological contribution to sarcopenia is thought to primarily occur through a loss of alpha motor neuron axons and the fiber type grouping that results from the concomitant remodeling of motor units: A decline in alpha motor neuron axons is observed in aging (Edström et al. 2007; McComas 1998) accompanied by changes in the motor neuron phenotype such as reduced dendritic tree size (Ramírez and Ulfhake 1992) and decreased synaptic input (Kullberg et al. 1998; Ramírez-León et al. 1999; Bergman and Ulfhake 2002). Surviving motor neurons compensate for the reduction in alpha motor neuron numbers by branching their axons to innervate the denervated muscle fibers (collateral sprouting), resulting in an increased number of muscle fibers per motor unit. However, it is unclear whether alteration of motor units is one of the first stages leading to sarcopenia.

To test our hypothesis that sarcopenia initiation and its early molecular programming could be detected by gene expression changes in nerves preceding sarcopenia development in muscle, we performed transcriptomic profiling of aging sciatic nerves *in vivo*. Signs of alterations in peripheral nerves prior to the clinical expression of sarcopenia is supported by a previous longitudinal study in C57BL/6J mice (aged between 4-24 months) that measured significant accumulation in the sciatic nerve of a series of proteins implicated in cytoskeleton dynamics/axonal transport and autophagic protein degradation pathways starting at 18 months (Krishnan et al. 2016), while in the same mice, loss of muscle mass and denervation at the NMJ is only clearly substantiated by 24 months of age (Barns et al. 2014; Chai et al. 2011). Another elegant study combining experiments in both rat and human tissues provided evidence that sarcopenia in vulnerable skeletal muscles of the lower limb (e.g., gastrocnemius) is tightly correlated with neuromuscular dysfunction (e.g., in the connected sciatic nerve), and that skeletal muscles resistant to sarcopenia (e.g., triceps brachii) are those with intact neuromuscular transmission (e.g., radial nerve) (Pannérec et al. 2016). In this study, while gene expression profiling in the muscles was studied at adult (8-10 months), early-sarcopenic (18-20 months), and confirmed sarcopenic (22-24 months) stages, in the linked peripheral nerves it was only examined in adult and old sarcopenic mice, limiting the opportunities to identify early molecular changes with a higher likelihood of driving the initiation and progression of sarcopenia.

The present study assessed gene expression in the sciatic nerve of mice aged 5, 18, 21, and 24 months using both untargeted RNA-seq and qRT-PCR, and measured expression of genes associated with muscle denervation in gastrocnemius muscle of the same mice using qRT-PCR. Altogether, we showed that expression of genes previously implicated in sarcopenia are differentially expressed (|LFC| > 1.0, FDR < 0.05) in nerve as early as 18 months compared to healthy young adult controls, namely *Myog* (for review see Riuzzi et al. 2018) which was up-regulated with a log_2_ fold change ~19 and *Arntl* (also known as *Bmal1*, Kondratov et al. 2006) which showed a nearly four-fold decrease. Pathway enrichment analysis revealed that up-regulated DEGs were borderline associated with the tight junction (*p* = 0.057) and AMPK signaling pathway (*p* = 0.08) while down-regulated DEGs were associated with biosynthesis and metabolic pathways. Both up- and down-regulated DEGs were associated with circadian rhythm, suggesting this is a particularly important pathway in aging sciatic nerve. We detected gene expression changes at 18 months, prior to the onset of sarcopenia, which we confirmed at 24 months by up-regulation of *Chrng, Chrnd, Runx1*, and *Gadd45a* in muscle by qRT-PCR, as upregulation of these genes has previously been demonstrated to coincide with sarcopenia onset (Barns et al. 2014).

To identify the earliest transcriptomic changes that could be occurring in sarcopenia, we closely examined gene expression in sciatic nerve of mice aged 18 months compared to mice aged 5 months. Mice aged 18-to 24-months are equivalent to 56-to 69-year old humans, and the presence of early signs of sarcopenia in C57BL/6J mice at ~20 months of age has been reported, demonstrated by significantly decreased grip strength, exercise endurance, muscle volume, and muscle mass of these mice compared to 10-week-old C57BL/6J mice (Kim and Hwang 2020). While most studies agree that 24- to 25-month-old mice represent a reliable animal model of sarcopenia, as both muscle mass and strength are affected at this stage and linked with NMJ loss (Barns et al. 2014), most studies utilizing the natural aging C57BL/6J mouse model of sarcopenia study mice older than 24 months of age (Xie et al. 2021), obscuring the study of molecular mechanisms early in the disease course. As early as 18 months, we detected upregulation of *Myog* in sciatic nerve which suggests disturbances in presynaptic terminals (NMJs) because expression of this gene increases strongly in denervated muscles. We also detected significant down-regulation of *Arntl* in 18-month-old mice. Mice deficient in brain and muscle *Arntl* have reduced lifespans and show symptoms of premature aging including sarcopenia (Kondratov et al. 2006). Thus, reduced *Arntl* expression in sciatic nerve at 18 months could indicate that pathways related to initiation of sarcopenia are unfolding at this time point.

*Arntl* is a core component of the circadian clock. We found dysregulation in other circadian clock genes, including period circadian regulator 2 and 3 (*Per2, Per3*) and *Npas2.* The importance of circadian rhythm maintenance for the structure, function, and metabolism of skeletal muscle is clear when observing the muscle phenotype in models of molecular clock disruption; cycle disruptions are linked to aging and the development of many chronic diseases including sarcopenia (Vitale et al. 2019). For example, circadian rhythm disruption was associated with an increased risk of sarcopenia in a population-based study in Korea which compared the risk of sarcopenia between night-shift workers and those who never worked night shifts (Choi et al. 2019). Our results suggest that impairment of circadian rhythm maintenance is an important factor in the development of sarcopenia that should be investigated further.

Additionally, our functional and pathway enrichment analysis of up-regulated DEGs identified the tight junction and AMPK signaling pathway as important pathways. The tight junction suggests the implication of the NMJ, while the appearance of the AMPK signaling pathway as an enriched pathway is a significant finding in the context of sarcopenia as this pathway plays a key role as a master regulator of cellular energy homeostasis (Mallick and Gupta 2021). The kinase is activated in response to stresses that deplete cellular ATP, such as low glucose or hypoxia. As a cellular energy sensor responding to low ATP levels, AMPK activation positively regulates signaling pathways that replenish cellular ATP supplies, including autophagy, and negatively regulates ATP-consuming biosynthetic processes such as protein synthesis. This agrees with our finding of down-regulation of biosynthetic and metabolic pathways. AMPK has also been implicated in a number of species as a critical modulator of aging through its interactions with mTOR and sirtuins, and thus this pathway could represent a relevant therapeutic target for the treatment of sarcopenia (Mallick and Gupta 2021).

We also performed functional enrichment analysis on clusters of genes with similar patterns of expression across age groups. We hypothesized that genes belonging to clusters 2 and 3, which showed progressive decreases and increases in expression over time, respectively, may be associated with natural aging. Indeed, these genes were related to regulation of transcription in agreement with previous reports of a systematic age-related decrease in regulation of transcription (Barns et al. 2014). These clusters were also associated with themes related to immune responses/responses to external stimuli, programmed cell death, and development and transcriptional regulation. Clusters 1 and 4 were of particular interest given their decrease and increase in expression at 18 months, respectively, which corresponds to the pre-onset of sarcopenia in C57BL/6J mice. Genes in cluster 1 were significantly associated with extracellular matrix organization and extracellular structure organization, suggestive of tissue remodeling with age. This is in agreement with previous reports in muscle of mice aged 24 months (Barns et al. 2014).

A limitation of this study is that we did not assess muscle strength/function or perform histology to validate the clinical and pathological onset and full development of sarcopenia and verify altered NMJ morphology and subsequent muscle denervation. However, these features of sarcopenia have previously been extensively characterized in the C57BL/6J mice we used for our study (Barns et al. 2014; Graber et al. 2015; Pannérec et al. 2016; Krishnan et al. 2016), and we confirmed that mice were sarcopenic by 24 months of age by replicating previous findings by Barns and colleagues demonstrating that expression of genes associated with NMJ denervation including *Chrnd, Runx1*, and *Gadd45a* are up-regulated in the muscle of mice aged 24 months. However, we did not detect significant up-regulation of *Myog*, although there was a nonsignificant increase in expression compared to expression in 5-month-old mice. Another limitation is that we performed RNA-seq on total RNA isolated from the entire sciatic nerve, as opposed to pure motor nerves. As is the case with many of the large nerves of the vertebrate nervous system, the sciatic nerve is a mixed-function nerve, meaning it is made up of the axons of both sensory and motor neurons. Therefore, in addition to motor neuron axons, we sequenced some sensory axons as well. Interestingly, the sympathetic nervous system has recently gained attention for its role in regulating skeletal muscle motor innervation as well as its potential implication in sarcopenia pathogenesis (Delbono et al. 2021). Glial cells are also in close contact with neurons within the sciatic nerve, and therefore in our study we measured some changes relevant to this other cellular compartment. Glial cells are instrumental in neural metabolism and regulation of neurotransmission; they were thus also proposed to potentially have a leading role in neuromuscular aging and sarcopenia (Kwan 2013). Future studies could use novel spatial imaging techniques such as the RNAScope™ to untangle the cellular origin of the key molecular changes we reported here.

In the current study, we investigated the mechanisms contributing to skeletal muscle weakness and atrophy during aging using an unbiased bioinformatics analysis. To our knowledge, this is the first report assessing the transcriptome of sciatic nerves in the context of sarcopenia using untargeted RNA-seq. The strengths of this study include the use of unbiased RNA-seq over microarrays, validation of our RNA-seq findings using qRT-PCR, and the use of multiple time points during the mouse life span with multiple (n=5-6) biological replicates. Our results suggest that pathophysiological processes contributing to sarcopenia may be unfolding early in the nerve and suggest that increased understanding of the key mechanisms regulating skeletal muscle denervation at the NMJ could be critical to identifying novel treatments and biomarkers for sarcopenia in older adults.

## 5. Acknowledgements

This work was supported by the National Institutes of Health (NIH) National Institute on Aging (NIA) Grant (R21AG052011) awarded to D.B.R. and S.K., the National Institute of Environmental Health Sciences (NIEHS) Grant ES009089, and by the Ruth L. Kirschstein National Research Service Award Individual Predoctoral Fellowship (F31ES030973, awarded to N.C.). S.C., H.J.R., D.A., and A.M. are supported by the European Union’s Horizon 2020 Research and Innovation Programme under the Marie Sklodowska-Curie grant agreement (No. 778003). This research was also funded in part through the National Cancer Institute (NCI) Cancer Center Support Grant (P30CA013696) and used the Genomics and High Throughput Screening Shared Resource. We thank Chaolin Zhang for his assistance in interpreting the RNA-seq data.

## 7. Ethical Statement

The present study was conducted in accordance with the Guiding Principles for the Care and Use of Laboratory Animals, as adopted by the IACUC of Columbia University (New York, New York, USA).

