## Supplemental Figures for "Transcriptomic analysis of aging mouse sciatic nerve reveals early pathways leading to sarcopenia"

**Supplemental Figure 1: Sample-level Quality Control with Principal Component Analysis (PCA) and Correlation Heatmap of Regularized-logarithm-transformed Normalized Gene Expression (N=24).**

PCA plot (A) with all samples (N=24), labeled with sample identifier. Pairwise correlation heatmap (B) with all samples (N=24). Scale bar = Pearson correlation coefficient (*r*).

A

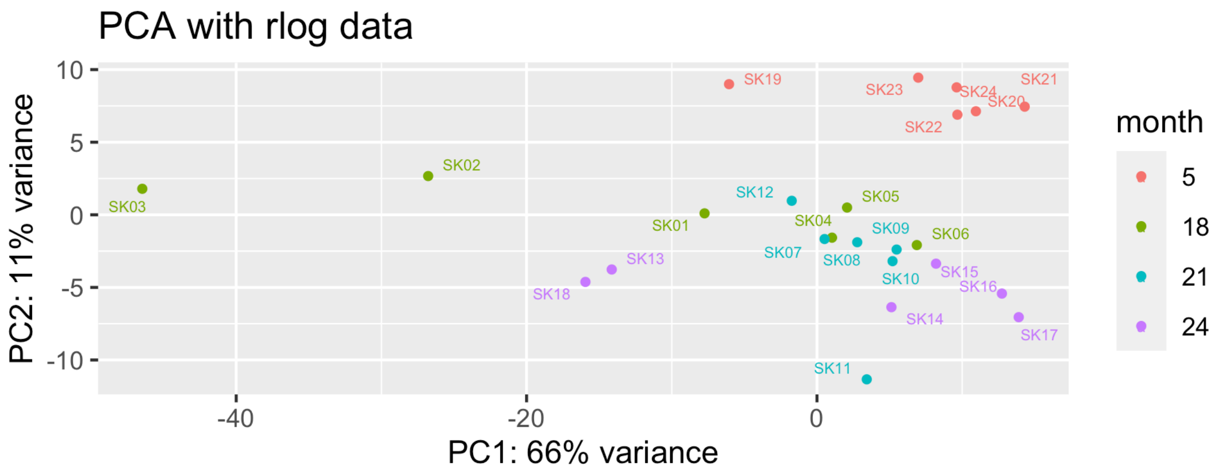


B

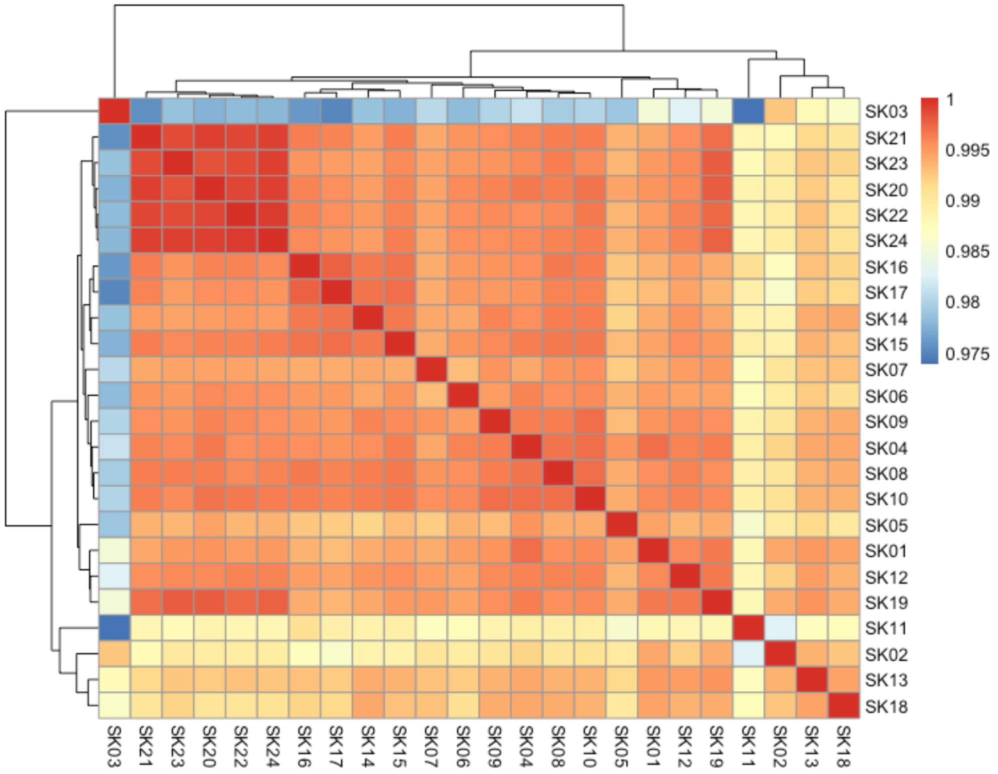

**Supplemental Figure 2: Final Principal Component Analysis (PCA) and Correlation Heatmap of Regularized-logarithm-transformed Normalized Gene Expression (N=22).** PCA plot (A) with final included samples (N=22), labeled with sample identifier. Pairwise correlation heatmap (B) with final included samples (N=22). Scale bar = Pearson correlation coefficient (*r*).

A

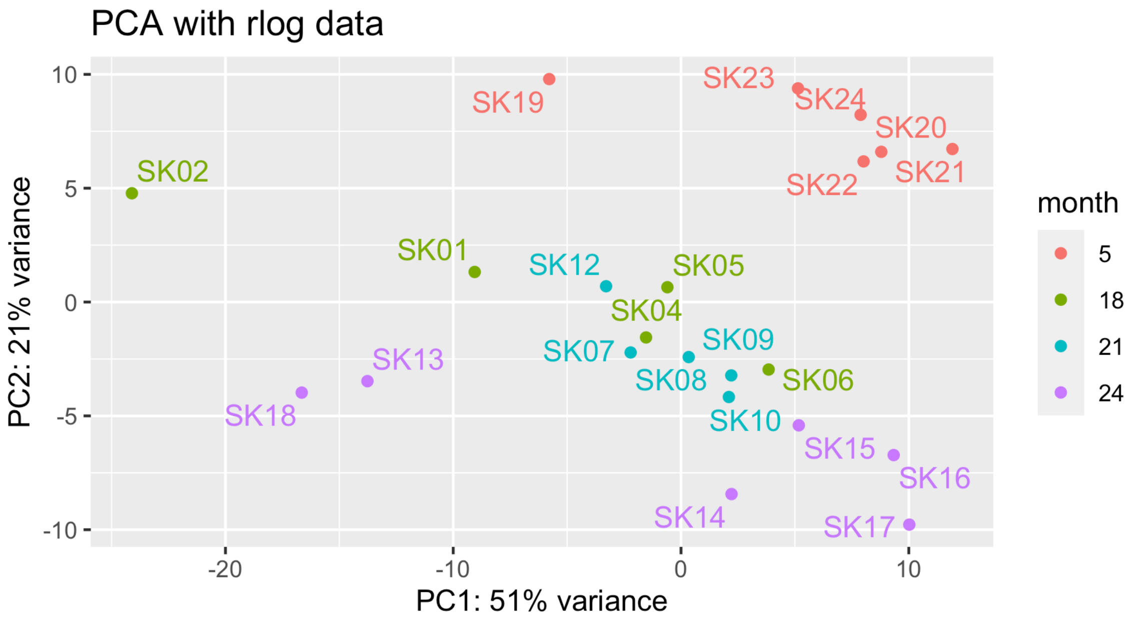

*Supplemental Figure 2 (continued).*

B

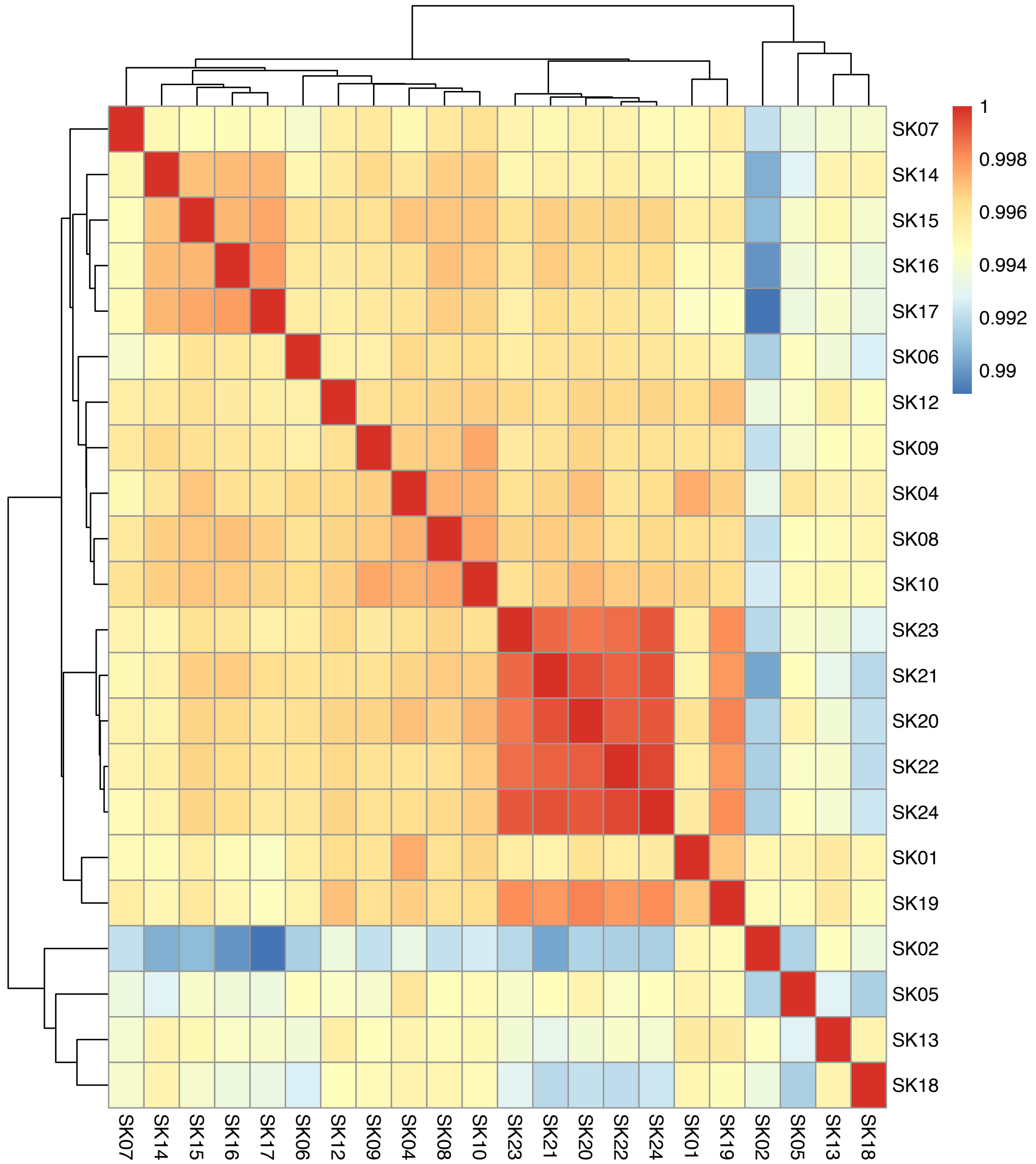

**Supplemental Figure 3: RT-PCR Cycle threshold (Ct) Values of Housekeeping Genes *Ppia* and *Hprt* in (A) Gastrocnemius muscle and (B) Sciatic nerve.** N=6 per group (except for *Hprt* measured in sciatic nerve, n=5 for 24-month-old age group).

A

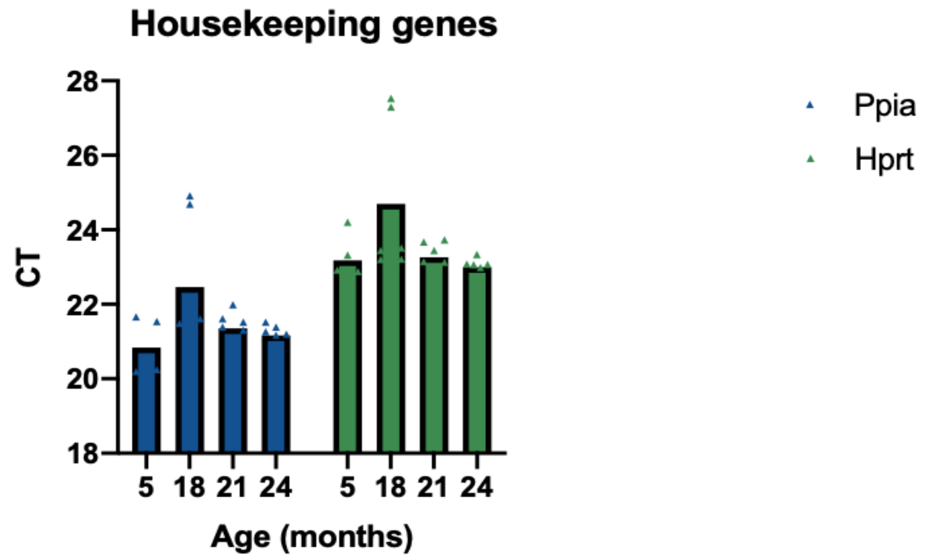


B

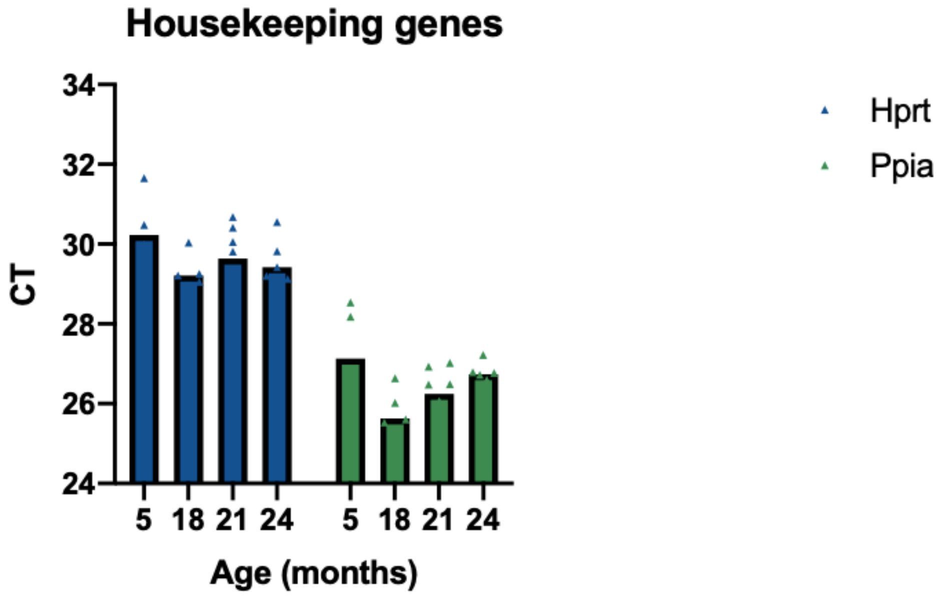

**Supplemental Figure 4. Expression of (A) *Chrng*, (B) *Chrnd*, (C) *Runx1*, (D) *Gadd45a*, and (E) *Myog* in Gastrocnemius Muscle of Mice Aged 18 to 24 months Compared to Mice Aged 5 Months.** Data are from 5-6 mice per group except for panel (A) which had 2-4 mice per group for mice aged 5-21 months as the other samples were below the Ct detection threshold. *significant difference between group with reference group at $p < 0.05$, ANOVA with Fisher's LSD tests. The delta-delta Ct was calculated using *Ppia* and *Hprt* housekeeping genes with 5-month-old mice used as the reference.

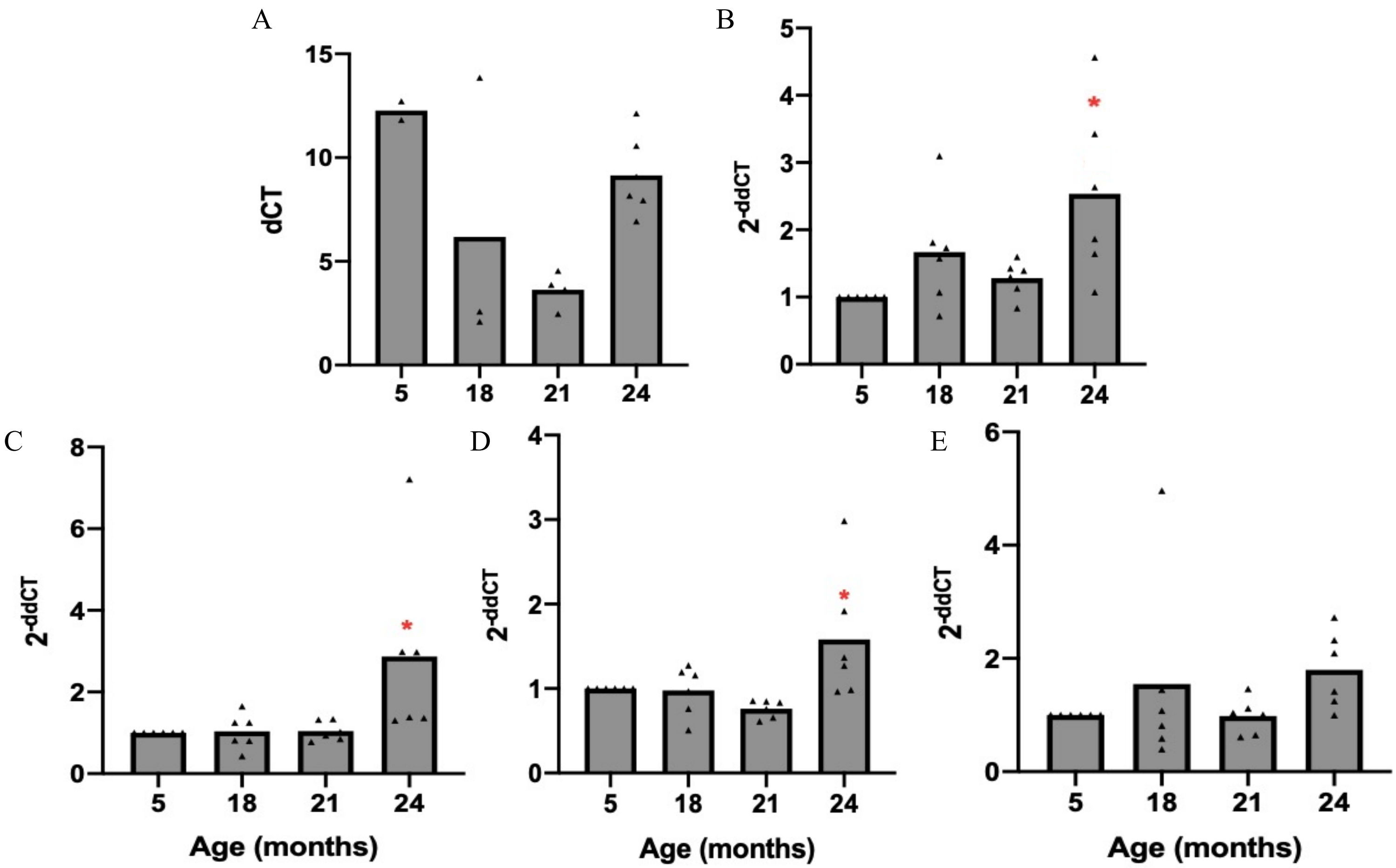

**Supplemental Figure 5: Volcano Plots of Differentially Expressed Genes (of 15,588 genes analyzed) in Sciatic Nerve Comparing Mice at Different Ages.**

Volcano plots of differentially expressed genes comparing (A) mice aged 24 months to 5 months; (B) mice aged 21 months to 5 months; (C) mice aged 21 months to 18 months; (D) mice aged 24 months to 18 months; (E) mice aged 24 months to 21 months. The dashed line is the FDR q-value cutoff of 0.05. Genes significant at |LFC| > 1.0 and FDR q-value < 0.05 are plotted in red. Sample sizes for groups of mice aged 5, 18, 21, and 24 months are n=6, 5, 5, and 6, respectively.

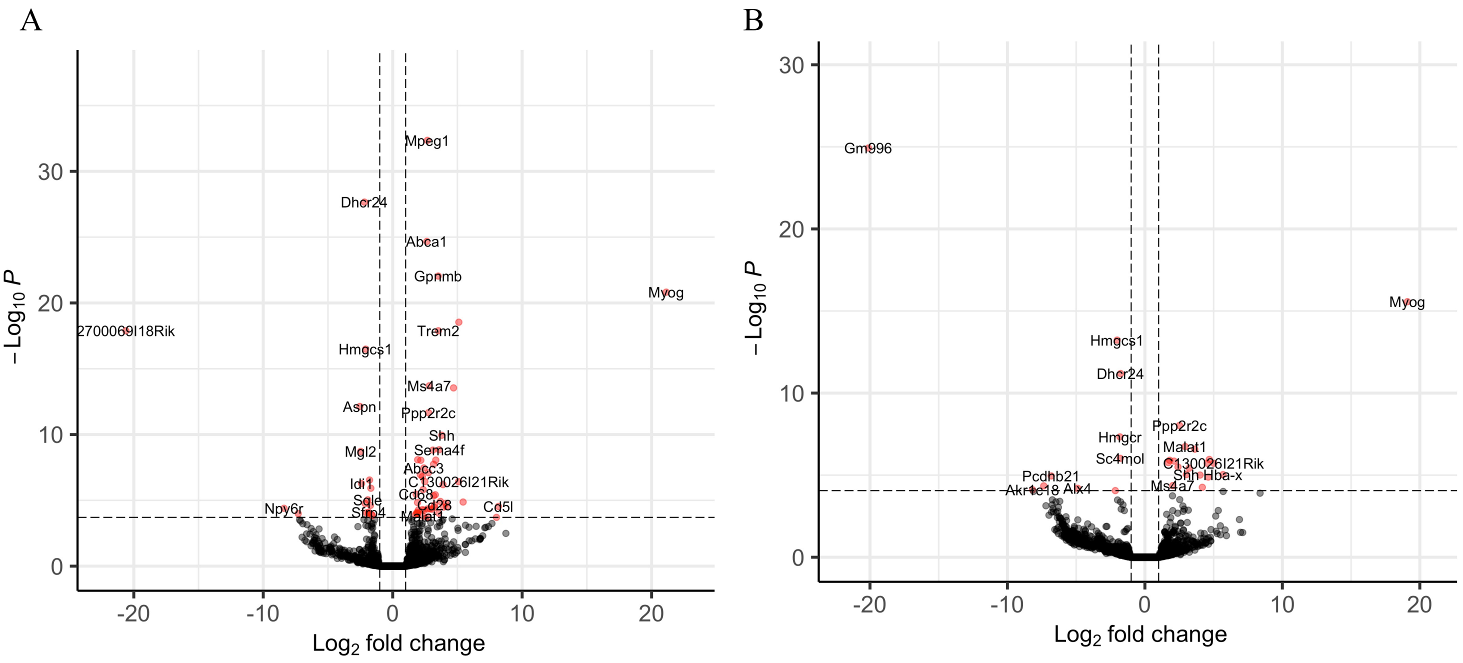

*Supplemental Figure 5 (continued).*

C

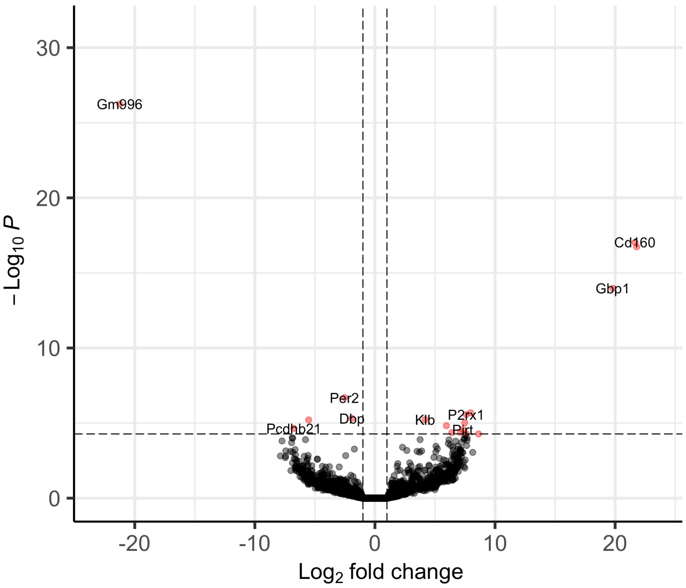


D

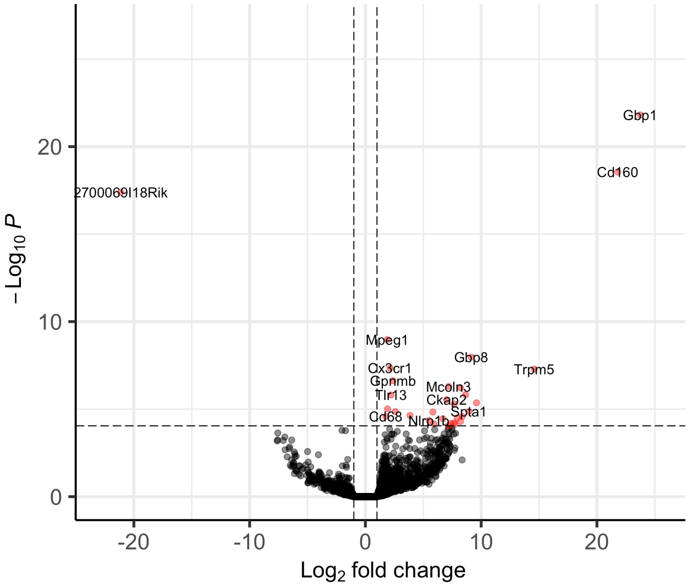


E

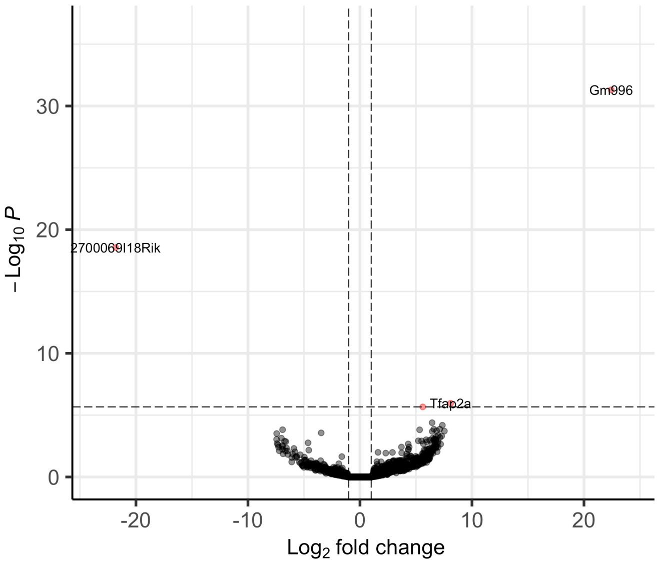

**Supplemental Figure 6. Expression of *Arntl* and *Npas2* in Sciatic Nerve of Mice Aged 5, 18, 21, and 24 months.** Plot of normalized DESeq2 counts of *Arntl* and *Npas2* in sciatic nerve of mice aged 5 (n=6), 18 (n=5), 21 (n=5), and 24 (n=6) months. *FDR *q*-value = 0.009, **FDR *q*-value = 0.0003, Wald test.

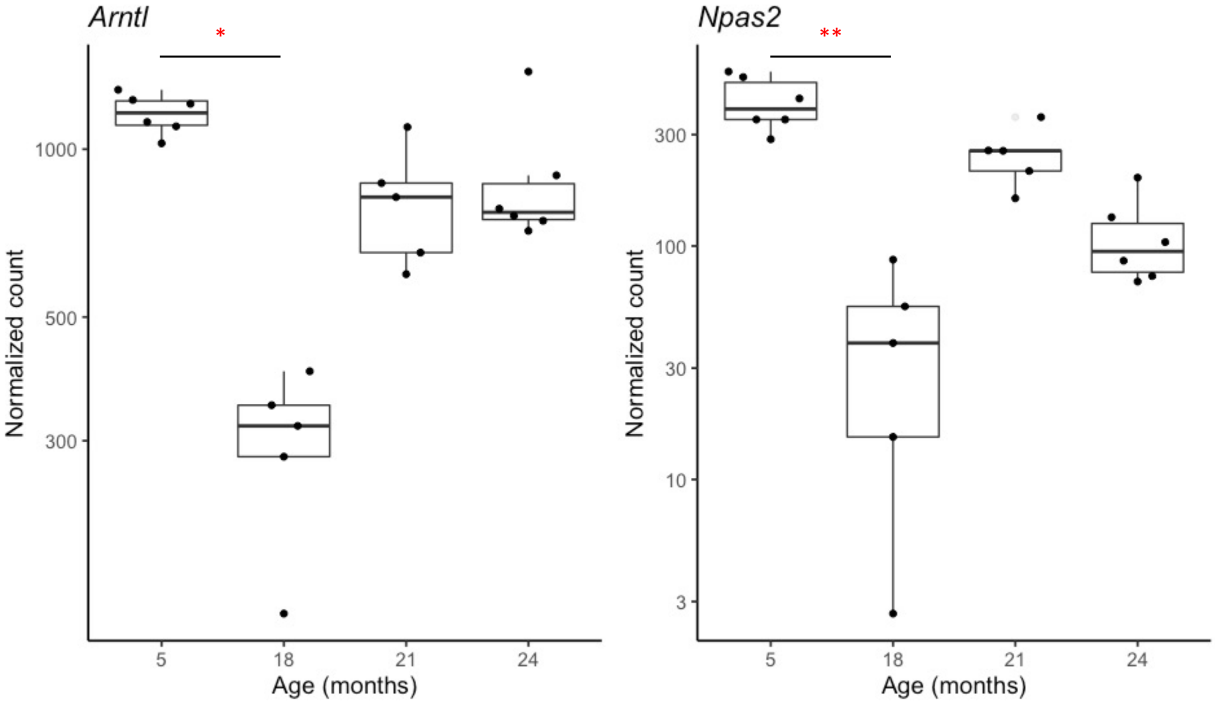
