## Supplemental Tables for "Transcriptomic analysis of aging mouse sciatic nerve reveals early pathways leading to sarcopenia"

| Sample ID | Age at death (months) | Sex | RNA Concentration (ng/µL) | Total RNA Submitted (ng) | Biolanalyzer RIN | RNA-seq read Alignment |
|---|---|---|---|---|---|---|
| SK01 | 18 | F | 3.1 | 310 | 7.9 | 59% |
| SK02 | 18 | F | 2.3 | 230 | 8.4 | 72% |
| SK03 | 18 | F | 4.9 | 490 | 8.9 | 67% |
| SK04 | 18 | F | 2.2 | 220 | 8.5 | 64% |
| SK05 | 18 | F | 5.2 | 520 | 6.2 | 50% |
| SK06 | 18 | F | 0.9 | 90 | 7.2 | 59% |
| SK07 | 21 | F | 3.5 | 350 | 9.3 | 60% |
| SK08 | 21 | F | 6.0 | 600 | 8.0 | 68% |
| SK09 | 21 | F | 3.0 | 300 | 8.9 | 70% |
| SK10 | 21 | F | 2.4 | 240 | 9.1 | 67% |
| SK11 | 21 | F | 3.9 | 390 | 8.8 | 71% |
| SK12 | 21 | F | 5.6 | 560 | 8.8 | 69% |
| SK13 | 24 | F | 9.4 | 940 | 8.7 | 60% |
| SK14 | 24 | F | 8.5 | 850 | 9.3 | 73% |
| SK15 | 24 | F | 6.2 | 620 | 8.6 | 69% |
| SK16 | 24 | F | 6.1 | 610 | 9.1 | 37% |
| SK17 | 24 | F | 9.2 | 920 | 9.0 | 66% |
| SK18 | 24 | F | 7.8 | 780 | 9.6 | 59% |
| SK19 | 5 | F | 2.3 | 230 | 8.1 | 82% |
| SK20 | 5 | F | 1.7 | 170 | 9.1 | 82% |
| SK21 | 5 | F | 2.0 | 200 | 9.4 | 80% |
| SK22 | 5 | F | 1.9 | 190 | 9.3 | 70% |
| SK23 | 5 | F | 1.9 | 190 | 9.4 | 73% |
| SK24 | 5 | F | 2.4 | 240 | 9.2 | 79% |

*Supplemental Table 1: Sample Information with RNA Quality Measures.*

Table includes sample identifiers, age at death (months), sex, RNA concentration (ng/µL), total RNA quantity (ng), RNA integrity number (RIN), and percent alignment to the genome.

F = Female, RIN = RNA Integrity Number (maximum = 10).

| Primer assay name | GeneGlobe ID | Catalog No. |
|---|---|---|
| Mm_Chrng_1_SG QuantiTect Primer Assay | QT00100268 | 249900 |
| Mm_Chrnd_1_SG QuantiTect Primer Assay | QT00199472 | 249900 |
| Mm_Ncam1_1_SG QuantiTect Primer Assay | QT00247695 | 249900 |
| Mm_Runx1_1_SG QuantiTect Primer Assay | QT00100380 | 249900 |
| Mm_Gadd45a_1_SG QuantiTect Primer Assay | QT00249655 | 249900 |
| Mm_Myog_1_SG QuantiTect Primer Assay | QT00112378 | 249900 |
| Mm_Hprt_1_SG QuantiTect Primer Assay | QT00166768 | 249900 |
| Mm_Ppia_1_SG QuantiTect Primer Assay | QT00247709 | 249900 |
| Mm_Dbp_1_SG QuantiTect Primer Assay | QT00103089 | 249900 |
| Mm_Ppp2r2c_1_SG QuantiTect Primer Assay | QT00122241 | 249900 |
| Mm_Per2_1_SG QuantiTect Primer Assay | QT00198366 | 249900 |
| Mm_Bhlhe41_1_SG QuantiTect Primer Assay | QT00141750 | 249900 |
| Mm_Cdh6_1_SG QuantiTect Primer Assay | QT01055159 | 249900 |
| Mm_Dhcr24_2_SG QuantiTect Primer Assay | QT01069712 | 249900 |
| Mm_Hmgcs1_1_SG QuantiTect Primer Assay | QT00132461 | 249900 |
| Mm_Col3a1_1_SG QuantiTect Primer Assay | QT01055516 | 249900 |
| Mm_Ngfr_1_SG QuantiTect Primer Assay | QT01047004 | 249900 |
| Mm_Traf2_1_SG QuantiTect Primer Assay | QT00103082 | 249900 |
| Mm_E2f1_1_SG QuantiTect Primer Assay | QT01079106 | 249900 |
| Mm_Mstn_1_SG QuantiTect Primer Assay | QT00093597 | 249900 |
| Mm_Mtor_1_SG QuantiTect Primer Assay | QT00118734 | 249900 |
| Mm_Fnip2_1_SG QuantiTect Primer Assay | QT00285544 | 249900 |
| Mm_Myo1f_2_SG QuantiTect Primer Assay | QT02329019 | 249900 |
| Mm_Bnip3_1_SG QuantiTect Primer Assay | QT00100233 | 249900 |
| Mm_Sod1_1_SG QuantiTect Primer Assay | QT00165039 | 249900 |
| Mm_Pink1_1_SG QuantiTect Primer Assay | QT00111349 | 249900 |
| Mm_Col11a1_1_SG QuantiTect Primer Assay | QT01055418 | 249900 |

*Supplemental Table 2: Qiagen QuantiTect Primer Assays with GeneGlobe ID.*

List of Qiagen primers used for qRT-PCR.

| | GO BP Term | Gene count | *P*-value | FDR *q*-value | Genes |
|---|---|---|---|---|---|
| Upregulated | Circadian rhythm | 4 | 5.70E-05 | 1.10E-02 | *Dbp, Lep, Per2, Per3* |
| | Negative regulation of transcription from RNA polymerase II promoter | 5 | 1.40E-03 | 1.30E-01 | *Hba-x, Hbb-y, Lep, Per2, Per3* |
| | Rhythmic process | 3 | 3.70E-03 | 2.30E-01 | *Dbp, Per3, Tef* |
| | Oxygen transport | 2 | 7.20E-03 | 3.40E-01 | *Hba-x, Hbb-y* |
| | Regulation of insulin secretion | 2 | 3.30E-02 | 8.60E-01 | *Lep, Per2* |
| | Multicellular organism development | 4 | 3.40E-02 | 8.60E-01 | *Dbp, Myog, Nnat, Tef* |
| | Muscle contraction | 2 | 3.50E-02 | 8.60E-01 | *Lmod2, Myh1* |
| | Transcription, DNA-templated | 5 | 3.90E-02 | 8.60E-01 | *Dbp, Myog, Per2, Per3, Tef* |
| | Circadian regulation of gene expression | 2 | 4.30E-02 | 8.60E-01 | *Per2, Per3* |
| | Cellular response to retinoic acid | 2 | 4.60E-02 | 8.60E-01 | *Lep, Myog* |
| Downregulated | Sterol biosynthetic process | 5 | 5.30E-09 | 9.40E-07 | *Dhcr24, Hmgcr, Hmgcs1, Mvd, Sqle* |
| | Cholesterol metabolic process | 5 | 7.20E-07 | 6.30E-05 | *Dhcr24, Hmgcr, Hmgcs1, Mvd, Sqle* |
| | Cholesterol biosynthetic process | 4 | 2.30E-06 | 1.30E-04 | *Dhcr24, Hmgcr, Hmgcs1, Mvd* |
| | Steroid biosynthetic process | 4 | 1.70E-05 | 7.50E-04 | *Dhcr24, Hmgcr, Hmgcs1, Mvd* |
| | Steroid metabolic process | 4 | 4.20E-05 | 1.50E-03 | *Dhcr24, Hmgcr, Hmgcs1, Mvd* |
| | Isoprenoid biosynthetic process | 3 | 8.70E-05 | 2.50E-03 | *Hmgcr, Hmgcs1, Mvd* |
| | Response to redox state | 2 | 5.80E-03 | 1.30E-01 | *Arntl, Npas2* |
| | Lipid metabolic process | 4 | 5.90E-03 | 1.30E-01 | *Dhcr24, Hmgcr, Hmgcs1, Mvd* |
| | Aging | 3 | 8.80E-03 | 1.70E-01 | *Hmgcr, Arg1, Col3a1* |

| | | | | | |
|---|---|---|---|---|---|
| | Response to vitamin E | 2 | 1.10E-02 | 1.90E-01 | *Hmgcs1, Arg1* |
| | Skin development | 2 | 4.70E-02 | 7.10E-01 | *Dhcr24, Col3a1* |
| | Circadian regulation of gene expression | 2 | 4.90E-02 | 7.10E-01 | *Arntl, Npas2* |

*Supplemental Table 3: GO functional enrichment analysis of up- and down-regulated DEGs comparing 18-month-old mice (n=5) to 5-month-old mice (n=6).* Shown are enriched GO BP terms with $p < 0.05$. BP: Biological Process.
