## Supplemental Spreadsheet Legends for "Transcriptomic analysis of aging mouse sciatic nerve reveals early pathways leading to sarcopenia"

*Supplemental Spreadsheet 1: DESeq2 results for all pairwise comparisons.*

Results for each pairwise comparison shown separately in each sheet, labeled by contrast.

baseMean: the base mean expression over all rows (average of the normalized count values, dividing by size factors, taken over all samples); lfcSE: log2 fold change standard error; padj: Benjamini-Hochberg adjusted *p*-value.

*Supplemental Spreadsheet 2: GOrilla functional enrichment analysis of clusters of genes with similar expression patterns across groups.* N = 2,925 (total number of genes in the analysis); B = Number of genes associated with a GO term; n = Number of genes in the target gene list (i.e., the number of genes in a given cluster); b = Number of genes in the target gene list also associated with a GO term.
